# Roles for the long non-coding RNA *Pax6os1*/*PAX6-AS1* in pancreatic beta cell identity and function

**DOI:** 10.1101/2020.07.17.209015

**Authors:** Livia Lopez-Noriega, Rebecca Callingham, Aida Martinez-Sánchez, Sameena Nawaz, Grazia Pizza, Nejc Haberman, Nevena Cvetesic, Marie-Sophie Nguyen-Tu, Boris Lenhard, Piero Marchetti, Lorenzo Piemonti, Eelco de Koning, A. M. James Shapiro, Paul R. Johnson, Isabelle Leclerc, Benoit Hastoy, Benoit R. Gauthier, Timothy J. Pullen, Guy A. Rutter

## Abstract

**Aim/Hypothesis:** Long non-coding RNAs (lncRNAs) are emerging as crucial regulators of beta cell development and function. Here, we investigate roles for an antisense lncRNA expressed from the *Pax6* locus (annotated as *Pax6os1* in mice and *PAX6-AS1* in humans) in beta cell identity and functionality.

**Methods:** *Pax6os1* expression was silenced in MIN6 cells using siRNAs and changes in gene expression were determined by RNA sequencing or qRT-PCR. Mice inactivated for *Pax6os1* and human *PAX6-AS1*-null EndoC-βH1 cells, were generated using CRISPR/Cas9 technology. Human islets were infected with lentiviral vectors bearing a targeted shRNA or *PAX6-AS1*, which were used to silence or overexpress, respectively, the lncRNA. RNA sequencing or RT-qPCR were used to measure transcriptomic changes and RNA pulldown in mice and human cells followed by mass spectrometry/western blot were performed to explore RNA protein interactions.

**Results:** *Pax6os1/PAX6-AS1* expression was upregulated at high glucose concentrations in derived beta cell lines as well as in mouse and human islets, and in pancreatic islets isolated from mice fed a high fat diet (n=6, p=0.003) and patients with type 2 diabetes (n=11-5, p<0.01). Silencing or deletion of *Pax6os1*/*PAX6-AS1* in MIN6 or EndoC-βH1cells increased the expression of several β-cell signature genes, including *PDX1* and *INS*. Female, but not male, *Pax6os1* null mice fed a high fat diet showed slightly enhanced glucose clearance. ShRNA-mediated silencing of *PAX6-AS1* in human islets robustly increased *INS* mRNA, enhanced glucose-stimulated insulin secretion and calcium dynamics, while overexpression of the lncRNA exerted opposing effects. *Pax6os1/AS-1* interacted with histones H3 and H4 in mouse and human cells, indicating a possible role for this lncRNA in histone modifications in both species.

**Conclusions:** Increased expression of *PAX6-AS1* at high glucose levels may impair beta cell functionality and thus contribute to the development of type 2 diabetes. Thus, targeting *PAX6-AS1* may provide a promising strategy to enhance insulin secretion and improve glucose homeostasis in this disease.

**Research in context:** *What is already known about the subject?:* Long non-coding RNAs (lncRNAs) are crucial components of the pancreatic islet regulome, whose misexpression may contribute to the development of diabetes.

*What is the key question?:* Is the lncRNA *Pax6os1/PAX6-AS1* involved in beta cell functionality and type 2 diabetes?

*What are the new findings?:* The expression of *Pax6os1/PAX6-AS1* is upregulated in mice fed a high fat diet and in pancreatic islets from type 2 diabetes donors. Overexpression of *PAX6-AS1* in human pancreatic islets reduces insulin expression, glucose stimulated secretion and intracellular calcium dynamics. Silencing *PAX6-AS1* in human pancreatic islets upregulates insulin expression, enhances glucose stimulated insulin secretion and increases intracellular calcium dynamics.

*How may this impact the clinic in the foreseeable future?:* Understanding the genetic factors induced by high glucose/obesity involved in beta cell dysfunction is crucial for the development of new therapies to treat T2D.

## Introduction

Type 2 Diabetes (T2D) develops when beta cells within pancreatic islets no longer secrete sufficient insulin to lower circulating blood glucose levels, usually in the presence of insulin resistance^1^. However, in a subset of T2D patients, defective insulin secretion is observed despite near-normal insulin sensitivity^2^. Therefore, in all forms of the disease, changes in beta cell “identity” are thought to play an important role in functional impairment and the selective loss of glucose responsiveness^3^.

Loss of normal beta cell function is often characterized by decreased expression of insulin (*INS*) and of genes critical for glucose entry and metabolism^4,3^. These changes may be accompanied by increased expression of so-called “disallowed genes”, whose levels are unusually low in healthy beta cells compared to other cell types^5^. Furthermore, in several models of diabetes, the above changes are associated with decreased expression of transcription factors that are required to maintain a mature beta cell phenotype, including pancreatic duodenum homeobox-1 (*PDX1*) ^7^ and MAF BZIP Transcription Factor A (*MAFA*)^6, 7^. The transcription factor Pax6 regulates the expression of several genes involved in insulin processing and secretion, while repressing signature genes defining different endocrine cell lineages, such as ghrelin (Ghrl)^8–10^. As a result, Pax6 expression is key to maintaining beta cell identity and function. Embryonic deletion of Pax6 in the murine pancreas leads to a drastic reduction in the number of alpha and beta cells, resulting in the death of mutant mice at postnatal days 3-6 due to severe hyperglycaemia^11^. Conditional inactivation of Pax6 in adult mice leads to impaired beta cell function and glucose intolerance^11, 12^, demonstrating the continued importance of this gene in mature beta cells. Further highlighting the importance of this locus in diabetes, pancreatic Pax6 cis-regulatory elements that interact with the Pax6 promoter and neighbouring long non-coding RNAs, modulate the activity of pancreas-related transcription factors such as Pax4^13^. In humans, loss-of-function mutations in *PAX6* are associated with aniridia (iris hypoplasia) and T2D^14,15^.

Long non-coding RNAs (lncRNAs), defined as transcripts > 200 nucleotides in length that are not translated into proteins, are crucial components of the pancreatic islet regulome, whose misexpression may contribute to the development of T2D^16^. LncRNAs are expressed in a tissue/cell specific manner and more than 1,100 have been identified in human and murine pancreatic islets^17,18^. Furthermore, the expression of several of these is modulated by high glucose concentrations, suggesting that they may be involved in beta cell compensation in response to high insulin demand^19^. Interestingly, a number of beta cell-enriched lncRNAs are mapped to genetic loci in the proximity of beta cell signature genes, such as *PLUTO*-*PDX1* or *Paupar-Pax6*, and regulate their expression in *cis*^19, 20^.

In the current study, we sought to determine whether a lncRNA expressed from the *PAX6* locus, previously annotated as *Pax6 opposite strand 1* (*Pax6os1*) in mice and *PAX6 antisense 1* (*PAX6-AS1*) in humans^21^, might impact beta cell identity and/or function through the modulation of *Pax6* expression or by other mechanisms.

## Methods

### Animals

All animal procedures were performed with approval from the EU Directive 2010/63/EU for animal experiments. Pax6os1 null mice experiments were performed with the approval of the British Home Office under the UK Animal (Scientific Procedures) Act 1986 (Project License PPL PA03F7F07 to I.L.) and from the local ethical committee (Animal Welfare and Ethics Review Board, AWERB) at Imperial College London. Db/db mice procedures were approved by the Andalusian Regional Ministry of Agriculture, Fish, Water and Rural Development (#3/11/2021/171/A and ALURES #nts-es-414463) and performed in accordance with the Spanish law on animal use RD 53/2013. Animals were housed in individually ventilated cages and kept under controlled environmental conditions (12 hs-light–dark cycle, 23 ± 1 °C with 30–50% relative humidity). Mice were provided with standard rodent chow, unless stated otherwise, and sterilized tap water *ad libitum.* For the HFD study, a chow enriched with 58% Fat and 25% Sucrose diet (D12331, Research Diet, New Brunswick, NJ) was used. Db/db mice were injected intraperitoneally either with scrambled or antisense oligonucleotides (5 mg/kg) purchased from QIAGEN^22^.

### Metabolic tests

For intraperitoneal glucose tolerance tests (IPGTT) and insulin measurements, 8–12-week-old animals were fasted for 16 h prior to experiments and received an intraperitoneal injection of glucose (1 g/kg or 3 g/kg respectively of body weight). Blood glucose levels were determined by tail venepuncture using a glucose meter (Accuchek; Roche, Burgess Hill, U). Insulin was determined by ELISA (CrystalChem, 90080), according to the manufacturer’s instructions.

### Human islets

Human islets were cultured in RPMI-1640 (11879-020) supplemented with 5.5 mM glucose, 10% FBS, 1% penicillin/streptomycin and 0.25 μg/ml fungizone. Characteristics of the donors are given in Suppl. Table 1.

### Cell culture

MIN6 were cultured with in DMEM (Sigma-Aldrich) (25mM glucose) supplemented with 15% (v/v) foetal bovine serum (FBS) and 2 mM L-glutamine.

EndoC-βH1 cells were cultured in DMEM medium (Thermo Fisher, 31885023) (5.5 Mm glucose) supplemented with albumin from bovine serum fraction V (BSA) (Roche), 50 μM 2-mercaptoethanol, 10 mM nicotinamide (VWR), 5.5 μg/ml transferrin (Sigma-Aldrich), 6.7 ng/ml sodium selenite (Sigma-Aldrich) and 1% penicillin/streptomycin.

### Small interfering RNA *Pax6os1* in MIN6

MIN6 cells were transfected with a pool of three small interfering RNAs (siRNAs) targeting *Pax6os1* or three control siRNAs using Lipofectamine™ RNAiMAX, according to the manufacturer’s protocol.

### CRISPR-Cas9 gene editing in mice and humans and lentivirus production

Guide RNAs targeting Pax6os1/PAX6-AS1 gene in mice and humans were designed using http://crispr.mit.edu and are described in supplemental Table 2.

For the generation of Pax6os1 null mice, guide RNAs were cloned into a pX330-U6-Chimeric_BB-CBh-hSpCas9 plasmid (Addgene, #42230). Pronuclear injection of the gRNAs was performed by the MRC transgenics unit, Imperial College London. F0 compound homozygous males (carrying different mutations in the *Pax6os1* gene) were crossed with WT females C57BL/6.

For CRISPR-Cas9 in EndoC-βH1 cells, guide RNAs were cloned into lentiCRIPSR v2 (Addgene #52961), modify to expressed Cas9 under the RIP promoter. EndoC-βH1 cells were then infected with lentiviral particles that were previously generated by co-transfection of lentiCRIPR v2 together with packaging (psPAX2; Addgene, # 12260) and envelope (pMD2.G; Addgene, #12259) plasmids into HEK293T cells.

### Transduction of pancreatic islets

Human islets were transduced as previously described^23^ with minor modifications. Briefly, islets were incubated overnight at 37 °C after arrival, mildly trypsinized (3 min, 0.5X Trypsin) and infected with lentiviruses. Experiments were performed 48 h after transduction.

### Massive parallel RNASequencing (RNASeq)

For MIN6 cells, libraries were constructed using a NEBNext Ultra II Directional RNA Library Prep Kit for Illumina (NEB) and NeBNext Multiplex Adapters (NEB) used for adapter ligation. Size selection of libraries was performed with SPRIselect Beads (Beckman Coulter). The Imperial BRC Genomics Facility performed sequencing as 75 bp paired end reads on a HiSeq4000 according to Illumina specification.

For EndoC-βH1 cells, libraries were prepared using RIBO-ZERO PLUS, following the manufacturer’s protocol. CABIMER Genomics Facility performed sequencing as 75 bp paired end reads Illumina Novaseq 6000, according to Illumina specifications.

Reads were aligned to the mouse (GRCm38) or human (GRCh38.p14) genome using HiSat2 or STAR 2.7.10 and quantified featureCounts or htseq, respectively. Differential expression was analysed with Deseq2 and clusterprofiler was used to determine enriched KEGG pathways.

### Immunoblotting

The antibodies used in this study are described in Supplemental Table 3.

### Click-iT EdU (5-ethynyl-2’-deoxyuridine and Cell viability assay

Control and *PAX6-AS1* null cells were fixed with 100% methanol, labelled for EdU using the Click-iT EdU Alexa Fluor 488 HCS Assay according to manufacturer’s instructions and co-stained with insulin. For cell viability, cells were cultured for 30 min. in 1 ml of PBS with Calcein-AM (1 µl) (Molecular Probes) and propidium iodide (1 µl) (Sigma-Aldrich). Cells were imaged using a Nikon spinning disk microscope at x20 magnification and counted using ImageJ software. At least 1000 cells were counted per experiment.

### Glucose-stimulated insulin and glucagon secretion

MIN6 or EndoCB-H1 cells were incubated with KRBH buffer (120 mM NaCl, 5 mM KCl, 4 mM CaCl2, 4 mM MgCl_2_, 25 mM NaHCO_3_ and 0.2% BSA, saturated with 95% O_2_/5% CO_2_; pH 7.4) supplemented with 3- and 30-Mm glucose (MIN6) or 0.5- mM and 17- mM glucose (EndoC-βH1) for 1 h. Cells were lysed for total insulin content with 1M Tris pH 8.0, 1% Triton, 10% glycerol, 5M NaCl and 0.2 EGTA.

For insulin secretion, groups of 15 islets were incubated for 30 min. in fresh KRBH buffer with 3 mM glucose at 37 °C under rotation in a water bath. After mix, briefly centrifuge (700 rpm, 2 min) and removing the supernatant, islets were further incubated with 500 ul of KRBH supplemented with 17 mM glucose. For glucagon secretion, groups of 15 islets were pre-incubated in e pre-incubated in KRB solution at 5.5 mM glucose twice for 20 min followed by incubation in 1 mM glucose^24^. Both total insulin and glucagon were extracted by adding acidified ethanol (75% ethanol/1.5% HCl). Insulin was measured by using a Homogeneous Time Resolved Fluorescence (HTRF) insulin assay kit (Cisbio) in a PHERAstar reader (BMG Labtech), following the manufacturer’s instructions. Glucagon was measured using a Glucagon ELISA kit (10-1271-01) from Mercodia according to manufacturer’s instructions.

### Intracellular free [Ca^2+^] measurements in islets

Groups of 15 islets were incubated with the Ca^2+^ indicator Cal-520 (Abcam, ab171868) in KRBH supplemented with 3 mM glucose for 45 min and imaged using a Zeiss Axiovert microscope equipped with a[×10 0.3–0.5 NA objective at 3mM and 17mM glucose as well as at 20 mM KCl.

### qRT-PCR

Total RNA was extracted using TRIZOL (Invitrogen, 15596026) or PureLink RNA mini kit (Invitrogen, 12183020) and on-column PureLink DNAse (Invitrogen, 12185010) following the manufacturer’s instructions. Complementary DNA (cDNA) was synthesized using random primers (Roche) and the High-Capacity cDNA Reverse Transcription kit (Life Technologies). Real-time qPCR was performed with a SYBR Green PCR master mix (Applied Biosystems). Primers used in this study are outlined in supplemental Table 4.

### Electrophysiology

Measurements were performed at 32°C in standard whole cell configuration using an EPC-10 amplifier and Pulse software. Exocytosis was measured using the increase of membrane capacitance in response to 500 ms depolarisations at 1 Hz. The extracellular medium was composed of: 118 NaCl mM, 5.6 KCl, mM, 2.6 CaCl mM2, 1.2 MgCl2, 5 HEPES, and 20 tetraethylammonium (TEA) (pH 7.4 with NaOH). The intracellular medium contained (in mM): 129 CsOH, 125 Glutamic acid, 20 CsCl, 15 NaCl, 1 MgCl2, 0.05 EGTA, 3 ATP, 0.1 cAMP, 5 HEPES (pH7.2 with CsOH). Cell size was estimated from the initial membrane capacitance before any stimulation. Calcium and sodium currents were measured from −60 mV to +50 mV and triggered by a 100 ms depolarisation from the resting potential (−70 mV). The effect of PAX6 loss of function on the depolarising inward current was measured at 3 independent time points: i) at the peak current occurring within the first 2ms is likely supported mainly by the rapidly inactivating voltage-gated sodium currents, ii) at 5ms of the depolarisation when the sodium component should be inactivated, iii) from 10-95 ms after the onset of the depolarization was used to determine the mean sustained current amplitude. The exocytosis and current measurements were normalised to the size of the cells reflecting the exocytosis and current densities expressed respectively in fF.pF^−1^ and pA.pF^−1^.

### MTT test

MTT activity was determined using the Cell Proliferation Kit I according to the recommendations of the manufacturer (Roche, Spain). Optical density was determined at 575 nm with a reference wavelength of 690 nm using a Varioskan Flash spectrophotometer (Thermo Scientific, Spain).

### mRNA stability assay

EndoC-βH1 cells were treated with 5 µg/ml of Actinomycin D (Sigma-Aldrich, A9415) for 4-, 8-, 24- and 32-hours prior RNA extraction^25^. Half-life of insulin at several time points was calculated using the online software https://calculator.academy/half-life-calculator/.

### Subcellular fractionation

MIN6 and EndoC-βH1 cells cells were lysed using 200 µl lysis buffer (10 mM NaCl, 2 mM MgCl, 10mM Hepes, 5mM dithiothreitol (DTT), 0.5% Igepal CA 630 (Sigma I3021)) and centrifuged at 8000 rpm, 4°C for 5 min. The supernatant was collected as the cytoplasmic fraction while the pellet was resuspended in 200 µl lysis buffer to yield the nuclear fraction.

### RNA pulldown

For RNA pulldown in MIN6, *Pax6os1* or *Slc16a1* (control) sequences were cloned using Sequence- and Ligation-Independent Cloning (SLIC) into a ptRNA-S1 plasmid that harbours a T7 promoter, *tRNA-S1 (encoding the streptavidin aptamer)* and a bovine growth hormone (BGH) polyadenylation site previously available in the lab. Primers for *Pax6os1* amplification are described in supplemental table 4. Two different SILAC MIN6 lysates were used with *Pax6os1* and *MCT1/SLC16A1* labelled with the heavy and light isotypes, respectively. Protein MIN6 lysates (4 mg) were incubated with streptavidin beads at 4°C overnight under rotation. After washes, the streptavidin aptamer and bound complex were eluted using 50 µl 10 mM Biotin (pH 7.2) (Sigma, B4501) suspended in the 1X Aptamer buffer and stored at −20°C. The pooled SILAC samples were run into an SDS-PAGE gel and cut into three slices. Each slice was subjected to in-gel tryptic digestion using a DigestPro automated digestion unit (Intavis Ltd.) and the resulting peptides were fractionated using an Ultimate 3000 nano-LC system in line with an Orbitrap Fusion Tribrid mass spectrometer (Thermo Scientific).

The RNA antisense pulldown in EndoC-βH1 cells was performed as described previously^26^. Briefly, 100,000,000 EndoC-βH1 were UV crosslinked at 254 for 0.8 J/cm^2^.Afterwards cells were lysed, sonicated and hybridized with DNA probes at 45 °C for 2:30 h with 1000 rpm intermittent mixing. The equivalent of 100,000 cells per group was used for the “RNA elution sample”, while the rest was used for the “Protein elution” fraction. Probes used in this study are described in Suppl. Table 5 and were purchased from IDT.

### Statistical analysis

For comparisons between two groups, statistical significance was calculated using non-paired two-tailed Student’s t-tests or paired student t-test for GSIS fold change in human islets, while repeated measurements two-way ANOVA tests were performed for metabolic tests in animals. All the statistical analyses were performed using Graph Pad Prism 8.0. or R version 4.1.1 (2021-08-10) and OriginPro2021b (OriginLab Corporation, Northampton, MA, USA). In all cases, a p-value < 0.05 was considered statistically significant. Error bars represent the standard error of the mean (SEM). Western blot analyses were performed using ImageJ software.

## Results

### *Pax6os1*/*PAX6-AS1* expression is enriched in pancreatic islets and upregulated by high glucose as well as in type 2 diabetes

The lncRNA *Pax6os1*/*PAX6-AS1* is a 1,464/1,656 nucleotide transcript mapped to a syntenically conserved region in chromosome 2 in mice and chromosome 11 in humans. It is transcribed antisense to the *Pax6* gene, overlapping with intron 1 in both species. The first intron of *Pax6os1*/*PAX6-AS1* also overlaps with *Paupar*, another lncRNA that is mainly expressed in α-cells and it is involved in Pax6 splicing^17^. As opposed to other lncRNAs including *Paupar*, *Pax6os1/PAX6-AS1* is not highly conserved at the nucleotide level between species, containing four exons in mice and three in humans, while important differences are also found in the predicted secondary structures (Figure 1A and 1B). The tissue distribution of *Pax6os1* in the mouse is similar to that of *Pax6*, being predominantly expressed in pancreatic islets and, to a lesser extent, in the eye and brain (Figure 1C). More specifically, within the islet, previously published data indicates that *Pax6os1* is enriched in β- and δ-cells ^27^, while there is no detectable expression in α-cells, where *Paupar* is strongly expressed (Suppl. Figure 1)^17^.

**Figure 1.**
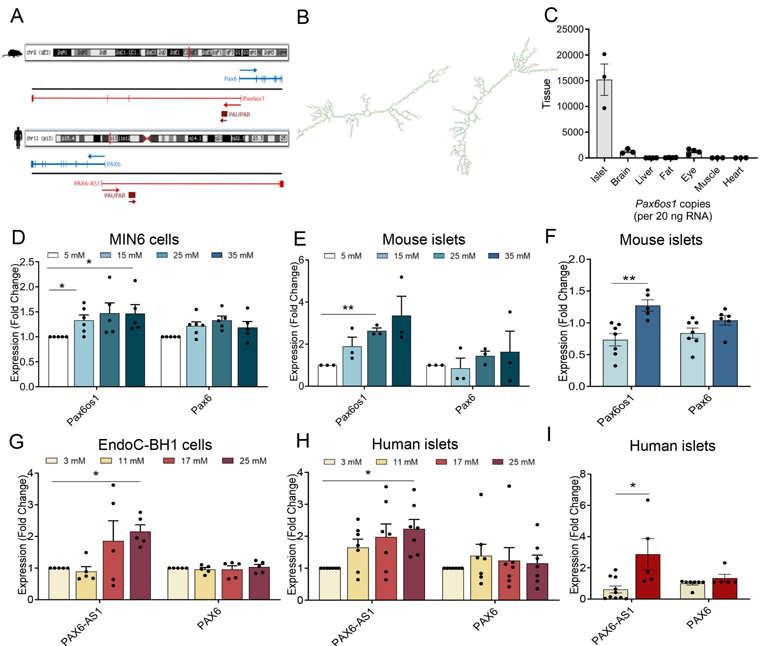
*Pax6os1* is chiefly expressed in pancreatic islets in the mouse and is upregulated by high fat diet as well as in islets from patients with type 2 diabetes. A) Schematic representation of the long non-coding RNA identified at the Pax6 locus in mice and humans. **B)** Secondary structure predicted using RNAfold from the VIENA package for Pax6os1 and PAX6-AS1, respectively and drew using FORNA webserver. **C)** Tissue distribution of *Pax6os1* expression. n=3. D, E) *Pax6os1* and Pax6 expression in MIN6 cells and CD1 mouse pancreatic islets cultured at different glucose concentrations for 48h (note that the standard glucose concentration for MIN6 cells culture is 25 mM). MIN6cells: n=6. CD1 islets: n= 3. F) *Pax6os1* and Pax6 expression in pancreatic islets from C57/BL6 mice in standard (STD) or high fat diet (HFD) for 8 weeks. n=6. G, H) *PAX6-AS1* and PAX6 mRNA expression in EndoC-βH1 cells and human islets cultured with different glucose concentrations for 48 hs (note that the standard concentration for EndoC-βH1 culture is 5.5 mM). EndoC-βH1 n=5. Human islets n=7. I) *PAX6-AS1* and PAX6 expression in human pancreatic islets from normoglycemic or diabetic donors. Control, n=10; Diabetic, n=5. Data are represented as the mean ± SEM. * p-value < 0.05 one-way ANOVA repeated measurements.

To determine whether *Pax6os1*/*PAX6-AS1* expression may be modulated under conditions of glucotoxicity, levels of the lncRNA were measured in both murine and human cell lines as well as in primary islets maintained at different glucose concentrations. Culture for 48 h in the presence of high glucose induced *Pax6os1* expression in both MIN6 cells (15 and 35 vs 5 mM glucose; *n*=5, p= 0.02 and 0.03, respectively) and CD1 mouse islets (11 vs 3 mM glucose; *n*=3, p<0.01) (Figure 1D-E). Furthermore, *Pax6os1* expression was increased in pancreatic islets from mice fed a high fat diet (HFD) compared to controls (*n*=6-5, p=0.003), while *Pax6* mRNA levels remained unaffected (Figure 1F). Likewise, *PAX6-AS1* expression was upregulated in the human Endoc-βH1 cell line (n=5, p=0.01) as well as in human pancreatic islets (n=7, p= 0.03) cultured at elevated glucose concentrations (Figure 1G-H). More importantly, expression of *PAX6-AS1* was substantially (4-5-fold) increased in islets from donors with T2D *versus* normoglycemic donors (*n*=11-5, p-value<0.01) (Figure 1I). In contrast, *PAX6* mRNA levels remained constant independently of the glucose concentration or disease status (Figure 1G-I).

### *Pax6os1* silencing upregulates beta cell signature genes in MIN6 cells

To explore the potential roles of *Pax6os1* in beta cell function or survival, we first transfected murine MIN6 cells with a small interfering RNA (siRNA) targeting the lnc-RNA. RNAseq analysis was then performed, and revealed that *Pax6os1* silencing (“knockdown”; KD) in MIN6 cells upregulated the expression of several beta cell signature genes, including *Ins1, Slc2a2 (Glut2) and Pax6* while further down-regulating several “disallowed genes” such as *Slc16a1* and *Ldha* (Figure 2A). In addition, enriched KEGG pathways in *Pax6os1* silenced MIN6 cells included “insulin secretion”, “Maturity onset diabetes of the young” and “Type II diabetes mellitus” (Figure 2B). qRT-PCR analyses cells confirmed a 35±5 % decrease in *Pax6os1* expression (p= 0.0005) as well as an increase in *Pax6* (1.28 ± 0.046-fold change; p=0.05), *Glut2*/*Slc2a2* (1.52 ± 0.15-fold change; p= 0.0144) and *Mafa* (1.72 ± 0.31-fold change; p=0.040) mRNA levels (Figure 2C). However, despite the upregulation of several beta cell signature genes, glucose-stimulated insulin secretion (GSIS), was not affected by *Pax6os1* silencing (Figure 2D-E).

**Figure 2.**
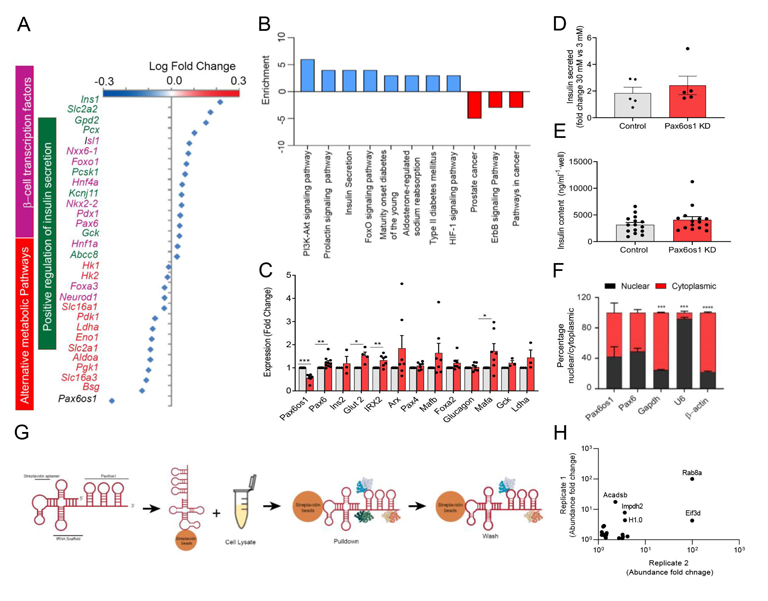
*Pax6os1* silencing upregulates beta cell signature genes in MIN6 cells. **A)** Differential expressed genes by *Pax6os1* knockdown as determined by RNA-seq performed in MIN6 cells 72 h post-transfection with s siRNA targeting *Pax6os1*. n= 4. **B)** KEGG pathway enrichment analysis relative to A. Significantly enriched KEGG pathways (p[<[0.05) are presented and the bar shows the fold-enrichment of the pathway. **C)** mRNA levels of β-cell signature genes and markers characteristic of other endocrine cell lineages in control and *Pax6os1* knockdown cells. n= 7. **D)** Fold change of insulin secreted relative to 3mM glucose. n= 5. **E)** Total insulin content. n = 14. **F)** Pax6os1 subcellular distribution. **G)** Schematic representation of Pax6os1 pulldown and MS. **H)** Relationship in abundance ratios above the 1.1 cut between the two experimental replicates performed. Top 5 hits are labelled. Short/branched chain-specific acyl-CoA dehydrogenase, mitochondrial 1 (Acadsb), Eukaryotic translation initiation factor 3 subunit D (Eif3d), Inosine-5’-monophosphate dehydrogenase 2 (Impdh2), Histone 1.0 (H1.0), Uncharacterized protein Rab8a (Rab8a). Data are represented as the mean ± SEM. *p-value< 0.05, student t-test.

We next sought to determine the mechanism by which *Pax6os1* may modulate the expression of target genes in MIN6 cells. LncRNAs may regulate gene expression through a number of different mechanisms, including chromatin remodelling, activation/repression of transcription factors in the nucleus as well as modulation of mRNA/protein stability in the cytoplasm^16^. Therefore, the subcellular localization of a lncRNA may provide a guide as to its mechanism(s) of action. Determinations of *Pax6os1* subcellular localization in MIN6 cells by subcellular fractionation indicated that the lncRNA was located in both the nucleus (∼40%) as well as in the cytoplasm (∼60%) (Figure 2F). Consistent with these results, both nuclear and cytoplasmic proteins were identified by mass spectrometry as binding protein partners of *Pax6os1* (Table 1). Interestingly, the top 5 hits included: Ras-related protein Rab8a, Eukaryotic translation initiation factor 3 subunit D (Eif3d), Inosine-5’-monophosphate dehydrogenase 2 (Impdh2), Short/branched chain-specific acyl-CoA dehydrogenase (Acadsb) and Histone H1.0 (Figure 2G-H). In addition, histones H4, H3.2, H2B, H1.1 and H1.4 in the nucleus as well as 3’-5’ RNA helicase YTHDC2 in the cytoplasm were also identified as *Pax6os1* binding partners (Table 1).

**Table 1.** Pax6os1 protein binding partners identified by mass spectrometry.

| Name | % Protein Coverage | Unique peptides | Average ratio (Control vs Pax6os1) | Cellular location |
| --- | --- | --- | --- | --- |
| Uncharacterized protein Rab8a | 16.35 | 1 | 100 | Nucleus |
| Eukaryotic translation initiation factor 3 subunit D | 1.64 | 1 | 52.13 | Cytoplasm |
| Histone H4 | 51.46 | 6 | 2.73 | Nucleus |
| Histone H1.0 | 23.71 | 4 | 4.03 | Nucleus |
| Inosine-5'-monophosphate dehydrogenase | 3.89 | 2 | 5.71 | Nucleus |
| Regulator of G-protein signalling 1 | 3.40 | 1 | 2.29 | Cytoplasm |
| Histone H3.2 | 18.23 | 4 | 2.38 | Nucleus |
| Histone H2B | 27.40 | 5 | 2.18 | Nucleus |
| Core histone macro-H2A.1 | 25.80 | 8 | 1.94 | Nucleus |
| Short/branched chain-specific acyl-CoA dehydrogenase, mitochondrial 1 | 4.95 | 2 | 9.93 | Mitochondria |
| Histone H1.1 | 32.86 | 7 | 1.45 | Nucleus |
| Pyrroline-5-carboxylate | 3.43 | 1 | 1.30 | Mitochondria |
| Uncharacterized protein (Fragment) Rpn2 | 2.12 | 1 | 1.36 | Endoplasmic reticulum |
| DEAD (Asp-Glu-Ala-Asp) box polypeptide 23 | 5.25 | 4 | 2.07 | Nucleus |
| 3'-5' RNA helicase YTHDC2 | 0.62 | 1 | 1.39 | Cytoplasm |
| Spectrin beta chain, non-erythrocytic 1 | 0.76 | 2 | 1.98 | Cytoskeleton |
| Histone H1.4 | 50.68 | 5 | 1.31 | Nucleus |
| Trifunctional enzyme subunit beta, mitochondrial | 8.42 | 4 | 1.48 | Mitochondria |

### *Pax6os1* deletion does not impact glucose homeostasis in T2D mouse models

In order to explore the possible consequences of *Pax6os1* loss for insulin secretion and glucose homeostasis *in vivo*, we used CRISPR/Cas9 gene editing to delete exon 1 of *Pax6os1* plus the immediate 5’ flanking region from the mouse genome in C57BL/6 mice (Figure 3A). Analysis of Super-Low Input Carrier- Cap analysis of gene expression (SLIC-CAGE) data (unpublished) in mouse islets, identified independent transcription start sites (TSS) for *Pax6* and *Pax6os1*, located ∼1kb apart (Suppl. Figure 2). Thus, the deletion generated spanned only the *Pax6os1* TSS and its putative promoter, as suggested by the presence of accessible chromatin in this region (ATAC-seq data), and of H3K4me3^28^ and H3K27Ac^29^ chromatin marks (Supplemental Figure 2A). Whereas *Pax6os1* expression was lowered by > 95 % in islets from KO mice, *Pax6* mRNA levels were unaffected (Figure 3B).

**Figure 3.**
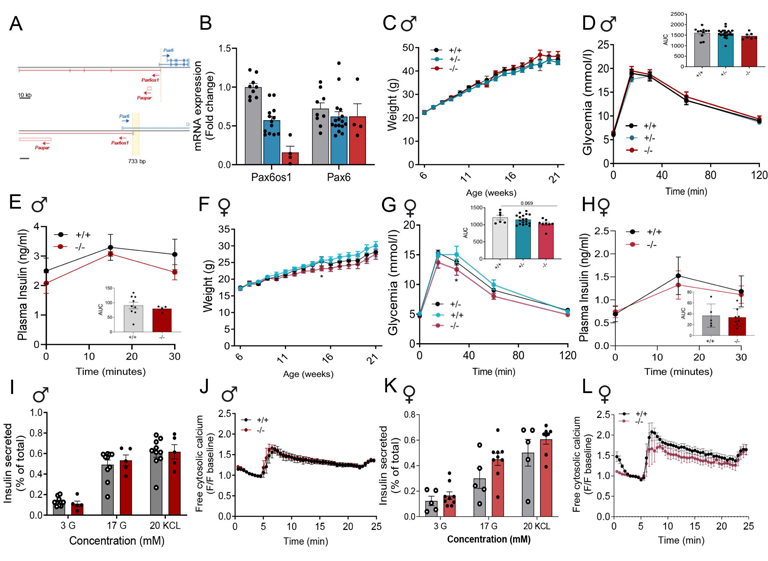
Pax6os1female null mice display mildly improved glucose tolerance and normal insulin secretion compared to WT animals under HFD. **A)** Schematic representation of the mutation generated in the Pax6os1 locus through CRISPR gene editing. B) *Pax6os1* and Pax6 expression in islets isolated from wt (+/+), *Pax6os1* heterozygous (+/-) and *Pax6os1* homozygous (-/-) mice. **C)** Body weights (g) of male Pax6os1 null mice under HFD. **D, E)** Circulating glucose levels and insulin in plasma after receiving an intraperitoneal load of glucose in male Pax6os1 null mice. Glucose: wt (n = 11), *Pax6os1* +/- (n = 22), *Pax6os1* -/-(n = 6); Insulin: wt (n= 9), *Pax6os1* (n=4). **F)** As panel C but in female mice. **G, H)** As panels D and E, respectively, but in female *Pax6os1* null mice. Glucose: wt (n=6), Pax6os1 +/- (n=18), Pax6os1-/-(n=8); Insulin wt(n=6), Pax6os1 -/-(n=8). **I)** Insulin secreted (represented as % of the total) at different glucose concentrations and after depolarization with KCl in pancreatic islets isolated from male Pax6os1 null mice. n= 10-5. **J)** Intracellular calcium in pancreatic islets isolated from male Pax6os1 null mice. n = 3.H, **K, L)** As panel I and J for female mice. n = 5-9 (I); n= 3 (J). Data are represented as the mean ± SEM. *p- value < 0.05 Repeated measurements two-way ANOVA.

No statistically significant differences were observed *in vivo* between wild-type (WT) and *Pax6os1* knockout (KO) male mice in weight, glucose clearance, insulin secretion under standard (STD) (Suppl Figure 3A-C) or HFD (Figure 3C-E). Similarly, no significant differences were observed between wild type or *Pax6os1* KO female mice in STD diet in weight, glucose clearance or insulin plasma levels (Supplemental Figure 3D-F). In contrast, *Pax6os1* KO female (but not male) mice under HFD displayed a tendency towards reduced body weight (Figure 3F) and significantly lower circulating glucose at 30 min. (p=0.041) during the IPGTT, with a strong trend towards a lower AUC during the experiment (WT: 1212±169 a.u. vs *Pax6os1* KO: 1030±134 a.u p=0.069) (Figure 3G). However, no differences were observed in insulin secretion *in vivo* (Figure 3H). Furthermore, there were no significant differences in GSIS or intracellular calcium dynamics between islets isolated from *Pax6os1* KO mice and wt independently of sex in STD (Suppl. 3-J) or HFD (Figure 3I-L).

In order to explore further the tendency observed in female mice in another animal model of type 2 diabetes, we treated *db/db* male and female mice with antisense oligonucleotides (ASO) targeting *Pax6os1*^22^. A significant and near-significant downregulation of *Pax6os1* in pancreatic islets could be observed after 4 weeks of treatment with ASO in female and male mice, respectively (Suppl Figure 4A-B). However, no significant differences were observed in body weight or glucose clearance between the different experimental groups (Suppl Figure 4C-H).

### *PAX6-AS1* depletion in EndoC-βH1 cells induces the expression of β-cell signature genes

In order to determine the effect of *PAX6-AS1* knockout in human beta cells, we used a tailored CRISPR/Cas9 approach to delete ∼80 bp within the first exon of *PAX6-AS1* from foetal human pancreas-derived EndoC-βH1 cells (Suppl Figure 5). Despite previously mentioned differences between *PAX6-AS1* and its ortholog in mouse, an RNA-seq analysis revealed several genes commonly modulated after silencing the lncRNA in MIN6 and EndoC-βH1 cells. Indeed, *PAX6-AS1*-depleted EndoC-βH1 cells displayed increased expression of the β-cell signature genes, *INS, PDX1* and *NKX6-1* (Figure 4A-B). However, no differences were observed in the expression of *LDHA* and *SLC16A1*, although the disallowed gene *SMAD3* was downregulated in *PAX6-AS1* KO compared to control cells. Further supporting the hypothesis that *PAX6-AS1* depletion favours the expression of β-cell signature genes over genes typically expressed in other endocrine cell types, SST expression was robustly reduced in *PAX6-AS1* KO cells (Figure 4A-B). Interestingly, although the calcium channels *CACNA2D1* and *CACNA2D2* were downregulated in *PAX6-AS1* depleted cells (Figure 4A-B), calcium signalling appeared as one of the KEGG pathways significantly activated after *PAX6-AS1* depletion due to the upregulation of RYR2 and FGF5 (Figure 4C-D). Other activated pathways included protein processing in the endoplasmic reticulum and the mitogenic secondary branch of insulin signalling Ras-MAPK (Figure 4C-D), while cAMP signalling and ECM-receptor interaction pathways were supressed (Figure 4C-D). Increased expression of *INS* (2.727 ± 0.6649-fold change, p=0.01) and *PDX1* (0.2625 ± 0.07348- foldchange, p-value = 0.01) was confirmed by qRT-PCR in PAX6-AS1 depleted cells (Figure 4E). Intriguingly, several genes that did not appeared to be modulated by *PAX6-AS1* depletion in our RNA-seq data, such as PAX6 (p-value = 0.009; p-adj = 0.15), showed increased mRNA levels in *PAX6-AS1* KO compared to control cells when measured by qRT-PCR (1.60 ± 0.16-fold change, p=0.002). However, no differences in *PAX6* protein levels were observed as measured by Western (immuno-) blot (Figure 4F-G), indicating that *PAX6* changes are unlikely to underly the effects of changes in *PAX6-AS1*.

**Figure 4.**
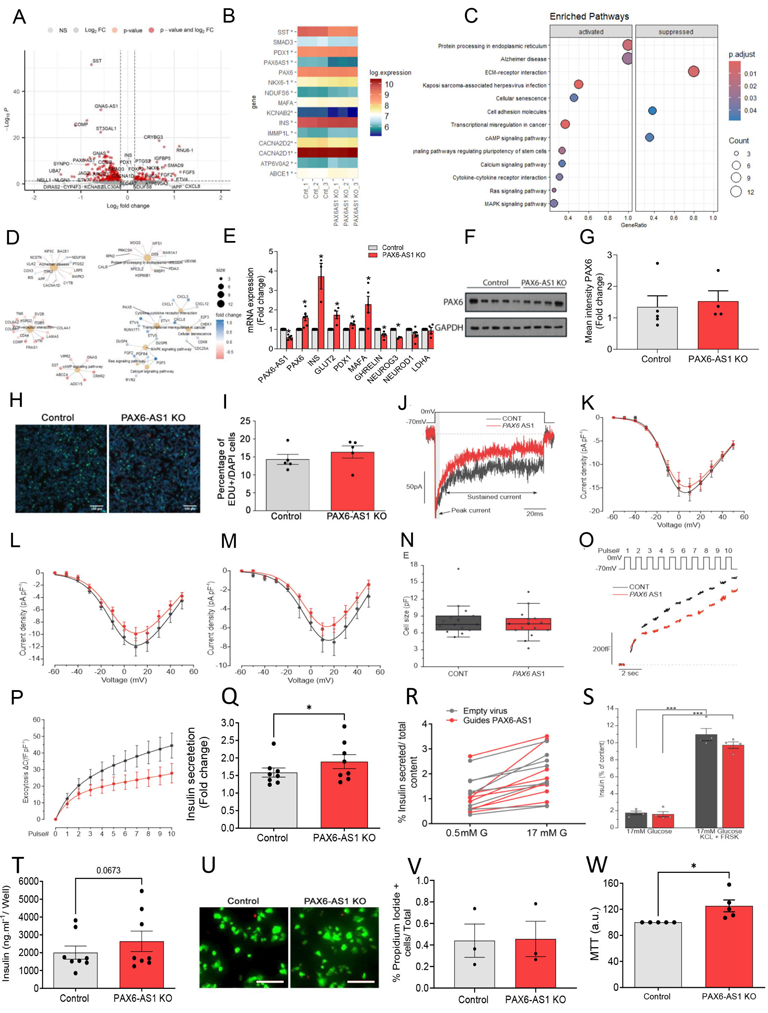
CRISPR/Cas9-mediated *PAX6-AS1* deletion from EndoC-βH1 cells increases insulin content without affecting cell number. **A)** Volcano plot representing genes significantly modulated in PAX6-AS1 depleted vs Control EndoC-βH1 cells. **B)** Heatmap representing the log expression for selected genes. **C, D)** Dotplot and cnetplot depicting the KEGG pathways significantly modulated in PAX6-AS1 KO EndoC-βH1 cells. **E)** mRNA expression of *PAX6-AS1*, beta cell signature genes and markers from other endocrine cell lineages in *PAX6-AS1* deleted EndoC-βH1 cells. *PAX6-AS1,* n= 6; *PAX6,* n= 6; *INS*, n= 4; *GLUT2/SLC2A2,* n= 5; *PDX1*, n= 4; GHRL, n= 4; NEUROG3, n= 3; NEUROD1, n= 5; LDHA, n=4. **F)** Western blot showing Pax6 protein levels. **G)** Densitometric analysis for F. **H, I)** Proliferation in PAX6-AS1 deleted EndoC-βH1 cells assessed by EdU staining: representative images (H) and quantification (I, n = 5). At least 1000 cells were counted per experiment using ImageJ software. **J)** Raw traces of mixed sodium and calcium currents elicited by a 100ms depolarisation from −70 to 0 mV in PAX6-AS1 KO and control cells. **K-M)** Quantification of the current density (see methods) at the peak which is mainly composed of the rapidly inactivating voltage-gated sodium and calcium currents (K), at 5ms when only the calcium component remains (the sodium component is inactivated) (L), and for the sustained component (M). n= 15. **N**) The cells measured in both conditions were of similar size. n= 15 cells in both cell types. **O)** Cumulative exocytosis was determined upon 10 depolarisations (pulses) from −70 to 0mV (top panel) using membrane capacitance measurement. **P)** Quantification of the cumulative exocytosis at each pulse. n=14-13. **Q, R)** Fold change of glucose induced insulin secretion. n= 8. **S)** Insulin secretion induced by 17 mM glucose and with addition of 35 mM KCl and 25 μM forskolin (FRSK). T) Determination of total insulin content per well. n= 8. **U, V)** Representative images showing calcein (green) and propidium iodide (red) staining and quantification of the percentage of propidium iodide positive cells. n = 3. At least 1000 cells were counted per experiment using ImageJ software. **W)** MTT assay. n = 5. Data are represented as the mean ± SEM. * p- value< 0.05, Student’s t-test or two-way ANOVA.

In spite of the activation of the mitogenic Ras-MAPK pathway, no differences were observed in the proliferation between control and *PAX6-AS1* KO EndoC-βH1 cells as determined by EdU staining (Figure 4H-I). Similarly, no significant differences were found in calcium currents between different cell types, which displayed similar cell size (Figure 4J-N). A strong tendency towards reduced exocytosis consistent with cAMP pathway suppression could also be observed in *PAX6-AS1* KO cells (Figure 4O-P). In contrast, *PAX6-AS1* depletion slightly enhanced GSIS in EndoC-βH1 cells as determined by an increased fold change in insulin secretion between 0.5 mM and 17 mM glucose (Control: 1.584 ±0.13, PAX6-AS1 KO: 1.894±0.19; p-value=0.041) (Figure 4Q-R), but not when cells were directly depolarized with KCl (Figure 4S).

Remarkably, the expression of SVB2-which is involved specifically in the exocytosis of GABA-containing synaptic-like microvesicles but not in insulin release^30,31^- was reduced in *PAX6-AS1* KO cells, while other genes involved in acidification and vesicular trafficking such as *ATPV0A2*^32,33^ were upregulated (Figure 4B). The improvement in GSIS was also accompanied by a tendency towards increased insulin content (Figure 4T) with no variations in cell number as suggested by the lack of significant differences in proliferation (Figure 4H-I) and cell death (Figure 4U-V). Furthermore, *PAX6-AS1*-depleted cells displayed increased mitochondrial activity as indicated by MTT assay (Figure 4W) and the upregulation of *NDUFS6* and *IMMP1L* (Figure 4B).

### *PAX6-AS1* directly interact with histones and modulate transcription of targe genes

In order to identify the molecular mechanisms of action of PAX6-AS1, we next sought to determine its subcellular localization in EndoC-βH1 cells. Remarkably, *PAX6-AS1* displayed the same expression pattern than *Pax6os1*, being located in the nucleus and the cytoplasm at similar proportions (∼40% and ∼60%, respectively) (Figure 5A). Next, we sought to validate in the human cell line the binding partners previously identified by mass spectrometry in MIN6 cells. To this end, we performed an RNA antisense pulldown (RAP), using biotinylated DNA probes antisense to our lncRNA followed by western blot analysis^26^ (Figure 5B). Successful RNA pulldown was confirmed by qRT-PCR in the RNA elution fraction, obtaining a [30% *PAX6-AS1* enrichment versus input using 4 specific probes targeting our lncRNA (Suppl table 5) (Figure 5C). In contrast, only [0.5% *PAX6-AS1* enrichment vs input was observed in the control group hybridized with probes targeting luciferase. Both groups showed <0.002% β*-ACTIN* enrichment versus input, confirming the specificity of our probes (Figure 5C). A direct interaction between *PAX6-AS1* and H3 as well as H4 was confirmed by western blot (Figure 5D), while direct binding of the lncRNA to other partners previously identified such as H1 or EIF3D could not be confirmed (data not shown). Consistent with these results, *PAX6-AS1* seemed to regulate *INS* expression at the transcriptional level as determined by increased levels of *INS* nascent mRNA and the lack of significant differences in mRNA stability between control (*INS* half-life: 11.29 ± 4.21 h) and PAX6-AS1 depleted cells (*INS* half-life: 9.04 ± 1.7 h) (p-value= 0.46) after treatment with actinomycin D (5 µg/ml) (Figure 5E-F).

**Figure 5.**
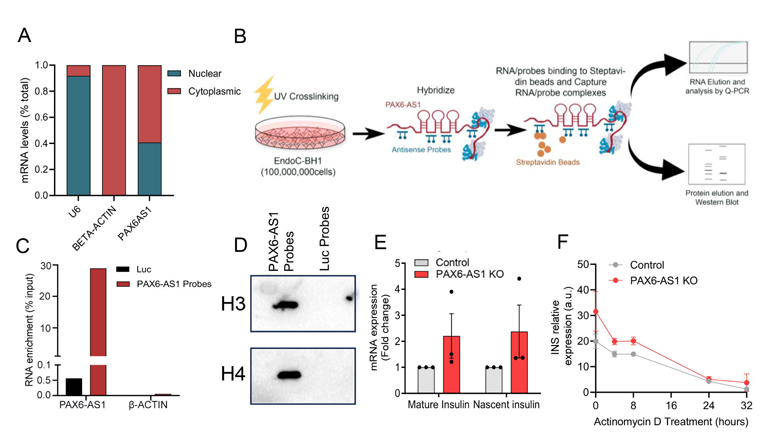
*PAX6-AS1* directly interacts to H3 and H4. **A)** Subcellular localization of PAX6- AS1 in EndoC-βH1 cells. **B)** Schematic representation of the RNA antisense pulldown. C) PAX-AS1 mRNA enrichment vs input in cells hybridized with probes targeting the lncRNA. **D)** Western blots showing H3 and H4 in cells hybridized with probes against our lncRNA or luciferase. **E)** Mature and nascent insulin mRNA expression as determined by Q-PCR in control and PAX6-AS1 depleted cells. n= 5. **F)** Determination of insulin mRNA stability as determined by Actinomycin D treatment in control and PAX6-AS1 KO cells. n= 6-3. Data are represented as the mean ± SEM. *p-value < 0.05, student’s t-test.

### *PAX6-AS1* knockdown enhances, whilst overexpression impairs, GSIS from human islets

To extend our results to fully differentiated human beta cells, we used lentiviral shRNA vectors to silence *PAX6-AS1* in pancreatic islets from post-mortem donors (Suppl Table 1). Transduced islets displayed a reduction in *PAX6-AS1* expression of 49 ± 12%, increased *INS* mRNA levels (2,727 ± 0,6649-fold change, *n*=5, p= 0.04), and reduced *GHRL* expression (0.57 ± 0.05-fold change, p< 0.0001) (Figure 6A). Importantly, the upregulation in INS mRNA levels was accompanied by enhanced GSIS (Scrambled: 3.44 ± 0.74-fold change vs *PAX6-AS1* shRNA: 6.69 ± 1.78-fold change, *n*=5, p=0.03), although total insulin content was not affected (Figure 6B-D). *PAX6-AS1*-silenced islets also showed increased intracellular Ca^2+^ dynamics in response to 17 mM glucose as assessed by the AUC for mean fluorescence (Scrambled: 13.09 ± 0.16 a.u. vs *PAX6-AS1* shRNA: 13.69 ± 0.10 a.u., p=0.049, paired t-test) (Figure 6E-F). In contrast, Ca^2+^ responses to plasma membrane depolarisation with KCl, added to open voltage-gated Ca^2+^ channels directly, were not significantly affected by *PAX6-AS1* silencing (Figure 6G). No additional effect was observed on glucagon secretion elicited by 1 mM glucose, suggesting that *PAX6-AS1* downregulation does not affect alpha cells (Figure 6H).

**Figure 6.**
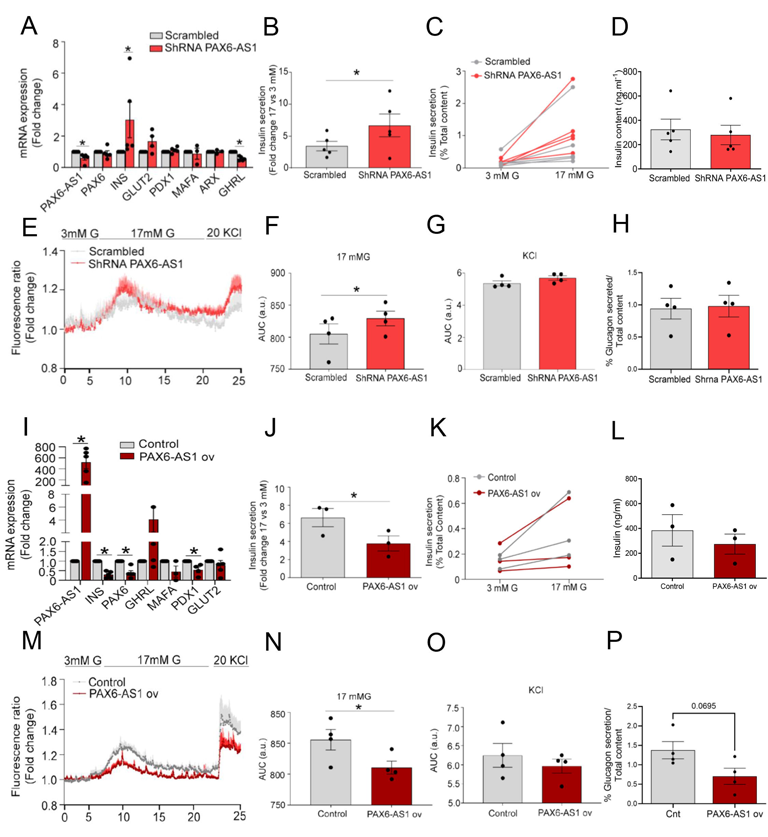
*PAX6-AS1* knockdown in human islets enhances glucose-induced insulin secretion, while *PAX6-AS1* overexpression has opposite effects. A) mRNA expression of *PAX6-AS1*, beta-signature genes and markers from other endocrine cell lineages in islets infected with a scrambled or a shRNA targeting PAX6-AS1. n= 4-5. **B, C)** Determination of pancreatic islet insulin content and glucose induced insulin secretion represented as the fold change in *PAX6-AS1* silenced islets. n= 5 (165,177,178, 182, 188). **D)** Trace showing calcium response in *PAX6-AS1* silenced islets. n= 4 (177,178, 182, 188). **E, F)** Area under the curve (AUC) for calcium dynamics in response to 17 mM glucose and 20 mM KCl, respectively in *PAX6-AS1* silenced islets. n=4. **G)** Glucagon secretion in *PAX6-AS1* silenced islets compared to scrambled. n=4 (R474, R480, R481, R4859). **H)** mRNA expression of PAX6-AS1 and beta cell signature genes in control and islets overexpressing PAX6-AS1. n= 6-3. **I, J)** Determination of pancreatic islet insulin content and glucose induced insulin secretion represented as the fold change in *PAX6-AS1* silenced islets. n= 3 (189, 190, 193). **K)** Trace showing calcium response in *PAX6-AS1* overexpressing islets. n= 4 (189, 190, 193, 196). **L, M)** Area under the curve (AUC) for calcium dynamics in response to 17 mM glucose and 20 mM KCl, respectively in *PAX6-AS1* overexpressing islets. n=4. **N)** Glucagon secretion in PAX6-AS1 overexpressing islets compared to control. n= 4 (R474, R480, R481, R4859). Data are represented as mean ± SEM. * p-value < 0.05, student’s t-test.

Demonstrating a deleterious effect on human beta cell function, *PAX6-AS1* overexpression in human islets led to a strong reduction in *INS* (0.29 ± 0.08-fold change, p< 0.0001) and *PDX1* (0.54 ± 0.14-fold change, p = 0.008) expression, while *GHR* was not affected (Figure 6I). The decrease in the expression of beta cell signature genes was accompanied by impaired GSIS (Control: 3.32 ± 0.5-fold change vs *PAX6-AS1* overexpression: 1.89 ± 0.41-fold change, p = 0.02), but unaltered total insulin content (Figure 6J-L). Correspondingly, *PAX6-AS1*-overexpressing islets displayed a significant reduction in intracellular Ca^2+^ dynamics in response to 17 mM glucose (Control: 16.05 ± 1.03 a.u. PAX6-AS1 overexpression: 15.01 ± 1.085, p=0.01, paired t-test) (Figure 6M-N), while there were no significant differences in the response to depolarisation with KCl (Figure 6O). In line with silencing experiments, overexpressing *PAX6-AS1* in islets did not significantly affect alpha cell functionality. Nevertheless, a tendency towards decreased glucagon secretion could be observed, suggesting that although this lncRNA is not normally expressed in alpha cells, forcing its expression can be detrimental in all pancreatic endocrine cells (Figure 6P).

## DISCUSSION

We show that *Pax6os1*/*PAX6-AS1*, a lncRNA transcribed from the *Pax6* locus and previously identified in the murine retina^18^, is chiefly expressed in beta cells within the pancreatic islet. This distribution differs from that of *Paupar*, also expressed from the *Pax6* locus, which is largely confined to α cells^17^. We demonstrate that *Pax6os1* is upregulated at high glucose concentrations, in an animal model of T2D (HFD) and in pancreatic islets from patients with this disease. Thus, it is tempting to speculate that increased expression of *PAX6-AS1* contributes to the pathogenesis of T2D.

Supporting this hypothesis, *Pax6os1* silencing increased the expression of several beta cell signature genes in murine MIN6 cells, while decreasing mRNA levels of disallowed genes, indicating a role for the lncRNA in beta cell identity and functionality. However, no significant differences were found in GSIS *in vitro*. Similarly, mice in which *Pax6os1* was deleted *in utero* displayed only a modest phenotype, which was evident in females maintained on a HFD diet.

Dissecting the functions of different transcripts within complex *loci* such as *Pax6/Pax6os1* is inherently challenging due to the close proximity of the transcriptional start sites. In this regard, it was conceivable that the DNA fragment deleted from the mouse genome (720 bp) by CRISPR/Cas9 to generate the *Pax6os1* knockout mouse might interfere with a regulatory region of *Pax6*, directly affecting the expression of the transcription factor. Indeed, assessment of the open chromatin state (by ATACSeq), and regulatory histone marks, indicated that the deletion of *Pax6os1* exon 1 and the proximal promoter region might potentially exert an effect on *Pax6* expression *in cis*. Furthermore, it is also possible that early developmental compensation occurred *in vivo* after *Pax6os1* deletion in the mouse, minimising the effects on β-cell function. Interestingly, however, the effects of *Pax6os1* inactivation were only observed in female mice, arguing against an effect mediated by alterations in Pax6 level (expected to affect both sexes equally).

In line with our results in MIN6 cells and in female mice, deletion of *PAX6-AS1* in the human beta cell line, EndoC-βH1, upregulated the expression of several beta-cell signature genes, including *Ins*. Furthermore, despite tendencies towards decrease calcium currents and decreased exocytosis, GSIS was slightly enhanced by *PAX6-AS1* deletion. This improvement in insulin release was accompanied by an apparent increase in mitochondrial activity and gene expression. Enhanced GSIS was also observed after lentivirus-mediated *PAX6-AS1* knockdown in human islets and this was accompanied by increased calcium dynamics in response to glucose. In contrast, islets overexpressing the lncRNA displayed impaired GSIS and cytosolic calcium dynamics. We note that *PAX6-AS1* is expressed in delta cells, albeit at lower levels than in beta cells, and that SST expression was strongly reduced in EndoC-βH1 cells. Therefore, impaired somatostatin secretion from delta cells after *PAX6-AS1* knockdown in islets may contribute to the direct effects of inactivating the lncRNA in the beta cell, further enhancing GSIS.

Since *PAX6-AS1* does not appear to modulate *PAX6* expression in humans, our hypothesis that *PAX6-AS1* exerts its major effects through the regulation of this transcription factor can be excluded. Interestingly, RNA pulldown experiments revealed that, amongst other partners, *Pax6os1/ PAX6-AS1* bind the histones H3 and H4, thus indicating that the lncRNA may regulate the transcription of target genes through epigenetic modifications.

LncRNAs have emerged in recent years as promising therapeutic targets in several diseases^16,34^. We show here that *PAX6-AS1* silencing in islets enhances insulin secretion, while increased expression of *PAX6-AS1-* as observed in T2D-may contribute to beta cell dysfunction and impaired GSIS. Although levels achieved in overexpression experiments were beyond the (patho-) physiological range, even in overexpressing cells, *PAX6-AS1* mRNA levels were still relatively low compared to most mRNAs and thus a non-specific toxic effect seems unlikely. Targeting *PAX6-AS1* might therefore provide a novel approach to maintain beta cell identity and functionality in T2D.

In summary, our data suggest that *Pax6os1/AS1* affects beta cell signature genes in both mice and humans, and that they share several binding partners. Nevertheless, there appear to be important species differences, with more marked effects observed in human islets and cell lines. Interestingly the effects of *Pax6os1* deletion were also sex-dependent in mice. Whether such differences also pertain in humans could not readily be explored here given the relatively small number of islet samples available and the use a cell line (EndoC-ßH1) from only one sex (female).

## Supporting information

All supplemental tables and figures

## Data availability

RNA sequencing (GEO) and proteomics (PRIDE) data will be uploaded and made publicly available at acceptance.

## Author contributions

L.L.N. and R.C. performed most of the experiments and analysed data. A.M-S contributed to experiments with human islets. G.P. prepared RNAseq libraries. A.M-S, N. H., N.C. and B.L. generated SLIC-CAGE and performed the analysis of ATAC-seq and histone marks in mouse islets. S.N. and B.H. performed electrophysiology. P.M., L.P., E.K., A.M.J.S. and P.J. provided human islets. M-S. N and I.L assisted with *in vivo* work. L.L.N. and G.A.R. wrote the manuscript. B.R.G., R.C. and A.M-S critically reviewed the manuscript. T.J.P. and G.A.R. conceived and supervised the study.

## Acknowledgments

G.A.R. was supported by Wellcome Trust Senior Investigator (WT098424AIA) and Investigator (WT212625/Z/18/Z) Awards, and MRC Programme grant (MR/R022259/1), Diabetes UK (BDA/11/0004210, BDA/15/0005275, BDA 16/0005485) and Imperial Confidence in Concept (ICiC) grants, an NIH-NIDDK project grant (R01DK135268) a CIHR-JDRF Team grant (CIHR-IRSC TDP-186358 and JDRF 4-SRA-2023-1182-S-N), CRCHUM start-up funds, and an Innovation Canada John R. Evans Leader Award (CFI 42649). G.A.R. and P.M. received funding from the European Union’s Horizon 2020 research and innovation programme via the Innovative Medicines Initiative 2 Joint Undertaking under grant agreement No 115881 (RHAPSODY). A.M-S was supported by an MRC New Investigator Research Grant (MR/P023223/1). B.H. was supported by RD Lawrence fellowship (BDA number:19/0005965). This project has received funding from the European Union’s Horizon 2020 research and innovation programme via the Innovative Medicines Initiative 2 Joint Undertaking under grant agreement No 115881 (RHAPSODY) to G.A.R and P.M. B.R.G. is supported by Ministerio de Ciencia E Innovación de España (PID2021-123083NB-I00 financed by MCIN/AEI/10.13039/501100011033 and by FEDER, UE) and the Juvenile Diabetes Research Foundation (3-SRA-2023-1307-S-B). L.L.N. is supported by the Consejería de Economía, Innovación y Ciencia (DOC_00652, FSE and Lema: Andalucía moves with Europe). We thank the Oxford Biomedical Research Centre (BRC), Diabetes Research and Wellness Foundation (DRWF), and Juvenile Diabetes Research Foundation (JDRF) for the provision of human islets.

## Twitter

@guy_rutter

## Twitter Statement

Long non-coding RNAs are emerging as important regulators of pancreatic beta cell function. Here we show that PAX6AS1/Pax6os1 plays an important role to control insulin secretion from both mouse and human beta cells and may provide a therapeutic target in some forms of type 2 diabetes.

## Abbreviations

EDU: 5-ethynyl-2’-deoxyuridine
GSIS: glucose-stimulated insulin secretion;
HbA1c: glycated haemoglobin
IPGTT: intraperitoneal glucose tolerance test
HFD: high fat diet
KO: Knockout
LncRNA: Long non-coding RNA
MTT: 3-(4,5-Dimethylthiazol-2-yl)-2,5-diphenyltetrazolium bromide
RNASeq: Massive parallel RNASequencing
siRNA: small interfering RNA
STD: Standard diet
T2D: Type 2 Diabetes

---

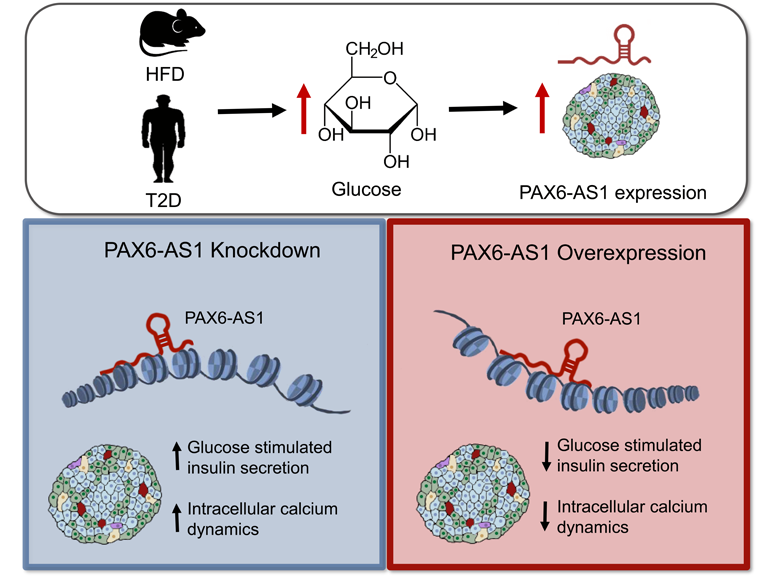

