## Supplementary material for "Roles for the long non-coding RNA *Pax6os1*/*PAX6-AS1* in pancreatic beta cell identity and function": All supplemental tables and figures

**Suppl. Table 1. List of donor characteristics and isolation centres.**

| **Identifier** | **Sex** | **Age** | **BMI** | **Isolation Centre** |
|---|---|---|---|---|
| 49 (T2D) | Male | 55 | 23.6 | Edmonton, Canada (Macdonald) |
| 60 | Male | 61 | 27.8 | Milan |
| 74 | Male | 83 | 24.5 | Pisa |
| 78 (T2D) | Female | 54 | 30.8 | Edmonton, Canada (McDonald) |
| 80 | Male | 54 | 35 | Edmonton, Canada (McDonald) |
| 85 | Female | 62 | 23.9 | Pisa |
| 91 (T2D) | Female | 53 | 21 | Leiden |
| 95 | Male | 38 | 42.6 | Edmonton, Canada (McDonald) |
| 101 (T2D) | Male | 57 | 35 | Leiden |
| 106 | Female | 49 | 20.57 | Milan |
| 114 | Female | 46 | 35 | Oxford |
| 116 | Female | 55 | 26 | Milan |
| 127 (T2D) | Male | 57 | 32 | Oxford |
| 165 | Female | 55 | 25 | Oxford |
| 177 | Male | 58 | 28.7 | Milan |
| 178 | Female | NA | 25.4 | Pisa |
| 182 | Male | 34 | 27 | Oxford |
| 188 | Male | 44 | 26 | Oxford |
| 189 | Female | 85 | 23.3 | Pisa |
| 190 | Female | 68 | 25.39 | Pisa |
| 193 | Male | 46 | 29.39 | Pisa |
| 196 | Male | 64 | 31.2 | Oxford |
| R474 | Male | 48 | NA | Edmonton, Canada (McDonald) |
| R480 | Female | 54 | NA | Edmonton, Canada (McDonald) |
| R481 | Female | 73 | NA | Edmonton, Canada (McDonald) |
| R485 | Male | 41 | NA | Edmonton, Canada (McDonald) |

**Suppl. Table 2. GuideRNAs used for CRISPR/Cas9 mediated disruption.**

| Human | Mouse |
|---|---|

| | |
|---|---|
| 5'-CACCGGTCCGGCCGCACGCCTTACC-3' | CACCGTGGTGGCCACTTTGCCCG |
| 5'-CACCGCAGGTCGCCTGCTTCGCAGT-3' | CACCGTTGTTTCCTCGGAGATCG |

**Suppl. Table 3. Antibodies used in this study**

| Antibody | Dilution | Vendor | Catalog number |
|---|---|---|---|
| Anti-Pax6 | 1:1000 WB, 1:100 IF | Biolegend | PRB-278P |
| Anti-GAPDH | 1:2000 | Cell Signaling | 2118 |
| Anti-H4 | 1:1000 | Cell Signaling | 2935 |
| Anti-H3 | 1:1000 | Sigma | H0164-25UL |
| Goat Anti-Rabbit (HRP) | 1:5000 | Abcam | ab6721 |
| Rabbit Anti-mouse (HRP) | 1:5000 | Sigma | A9044 |

**Suppl. Table 4. Primers used in this study.**

| Gene | Forward | Reverse |
|---|---|---|
| *Pax6os1-202* (mouse) | AGATGCCTTAGACAAGCCTG | ATTCACCTTCTTGGACCCTG |
| *Pax6os1-201* (mouse) | AGATGCCTTAGACAAGCCTG | ATTCACCTTCTTGGACCCTG |
| Pax6 (mouse) | ATGGGCGGAGTTATGATACCT | TGAAATGAGTCCTGTTGAAGTG |
| Beta Actin (mouse) | CGAGTCGCGTCCACCC | CATCCATGGCGAACTGGTG |
| Insulin 2 (mouse) | AGTAACCACCAGCCCTAAGTG | AGCACTGATCTACAATGCCAC |
| SLC2A2 (Glut 2) (mouse) | TTACAGTCACACCAGCATACAC | GCTTTGATCCTTCCAAGTTTGTC |
| Pdx1 (mouse) | GATGAAATCCACCAAAGCTC | TCGGTCAAGTTCAACATCAC |

| | | |
|---|---|---|
| Foxa2 (mouse) | CCCATTCTGGACATGGTGAAA | AGCACGCAGAAACCATAAATTAAA |
| Arx (mouse) | CCGCTGGGTCTGAGCACTT | GAAAAGAGCCTGCCAAATGC |
| Pax4 (mouse) | ATCCAGAACCAGTCCCAAAGAG | CCAACTGGCAAACTGAAAACG |
| Mafa (mouse) | CAGGTGGAGCAGCTGAAGCT | CCGCCAACTTCTCGTATTTCTC |
| Mafb (mouse) | CGCGTCCAGCAGAAACATC | AGCTGCTCCACCTGCTGAAT |
| Ghrelin (mouse) | GCTGGAGATCAGGTTCAATGC | CTGCTGATACTGAGCTCCTGACA |
| Irx2 (mouse) | GAGGACGAAGGGATCAGTCTACA | CGGCAGGGCAATTTTTCC |
| Gapdh (mouse) | AGGTCGGTGTGAACGGATTTG | GGGGTCGTTGATGGCAACA |
| Ldha | ATGAAGGACTTGGCGGATGA | ATCTCGCCCTTGAGTTTGTCTT |
| RNA, U6 small nuclear 1 (RNU6-1) (mouse) | CGATACAGAGAAGATTAGCATGG | AATATGGAACGCTTCACGA |
| Gck (mouse) | CAACTGGACCAAGGGCTTCAA | TGTGGCCACCGTGTCATTC |
| *PAX6-AS1* (human) | CAGCTCCAGGGAGAGGAAC | GAAGACACTCCTCCAGCAGAA |
| *PAX6-AS1* (human) | AGCTGCTGCCTTTCTCAAAA | CATTACTGCTGAGGGCCTTG |
| INS (human) | GCAGCCTTTGTGAACCAACA | ACCTGCCCCACCTGCAG |
| Intronic INS (human) | TTGATGACCGCAGATTCAAG | CCCCATCTCCTGACTATGGA |
| PAX6 (human) | CCGTGTGCCTCAACCGTA | CACGGTTTACTGGGTCTGG |
| Cyclophilin (human) | TATCTGCACTGCCAAGACTGA | CCACAATGCTCATGCCTTCTTTCA |
| MAFA (human) | GCCATCGAGTACGTCAACGA | CGGGAGGCTCCTTCTTCAC |
| MAFB (human) | TTCTTTGGGTGAGAAGGGATCGCA | TCAGCTTGCTGCCACGTTCTCTAT |

| | | |
|---|---|---|
| NEUROD1 (human) | ATTGCACCAGCCCTTCCTTTGATG | TCGCTGCAGGATAGTGCATGGTAA |
| NEUROG3 (human) | TAAGAGCGAGTTGGCACTGAGCAA | TTTGAGTCAGCGCCCAGATGTAGT |
| PDX1 (human) | TACTGGATTGGCGTTGTTTGTGGC | AGGGAGCCTTCCAATGTGTATGGT |
| SLC2A2 (human) | AGCTGCATTCAGCAATTGGACCTG | ATGTGAACAGGGTAAAGGCCAGGA |
| LDHA (human) | AGCCCGATTCCGTTACCT | CACCAGCAACATTCATTCCA |
| GHRL (human) | GGAAGATGGAGGTCAAGCAG | GCCTCTTCCCAGAGGATGTC |

**Suppl. Table 5. DNA probes used in this study.**

| | |
|---|---|
| **PAX6-AS1.1** | [Biotin~5]GGGCAGCTGGAGAGCTGGTGCTGTGGGGAACTGCACCATTAGTCCTTCCTG |
| **PAX6-AS1.2** | [Biotin~5]AATGGTACAAGCATAGAGCCAAACTCTGCCCTCTGCGAGGTGCTGCTCCCA |
| **PAX6-AS1.3** | [Biotin~5]TTAGACTCGTAAGCCATTAAATATTAAGACTTTGTCAACGTGGCCCTGTGCACT |
| **PAX6-AS1.3** | [Biotin~5]CTGGGACAGACATGAACGTGTCTGTCTCTCCAGAAGATATTCAGCTTCGTGT |
| **Luc-1** | [Biotin~5]TCCATCCTCTAGAGGATAGAATGGCGCCGGGCCTTTCTTTATGTTTTTGGCGTCTTCCAT |
| **Luc-2** | [Biotin~5]CCCTTAGGTAACCCAGTAGACCCAGAGGAATTCATTATCAGTGCAATTGTTTTGTCACGA |
| **Luc-3** | [Biotin~5]TGTTGGGGTGTTGTAACAATATCGATTCCAATTCAGCGGGGGCCACCTGATATCCTTTGT |

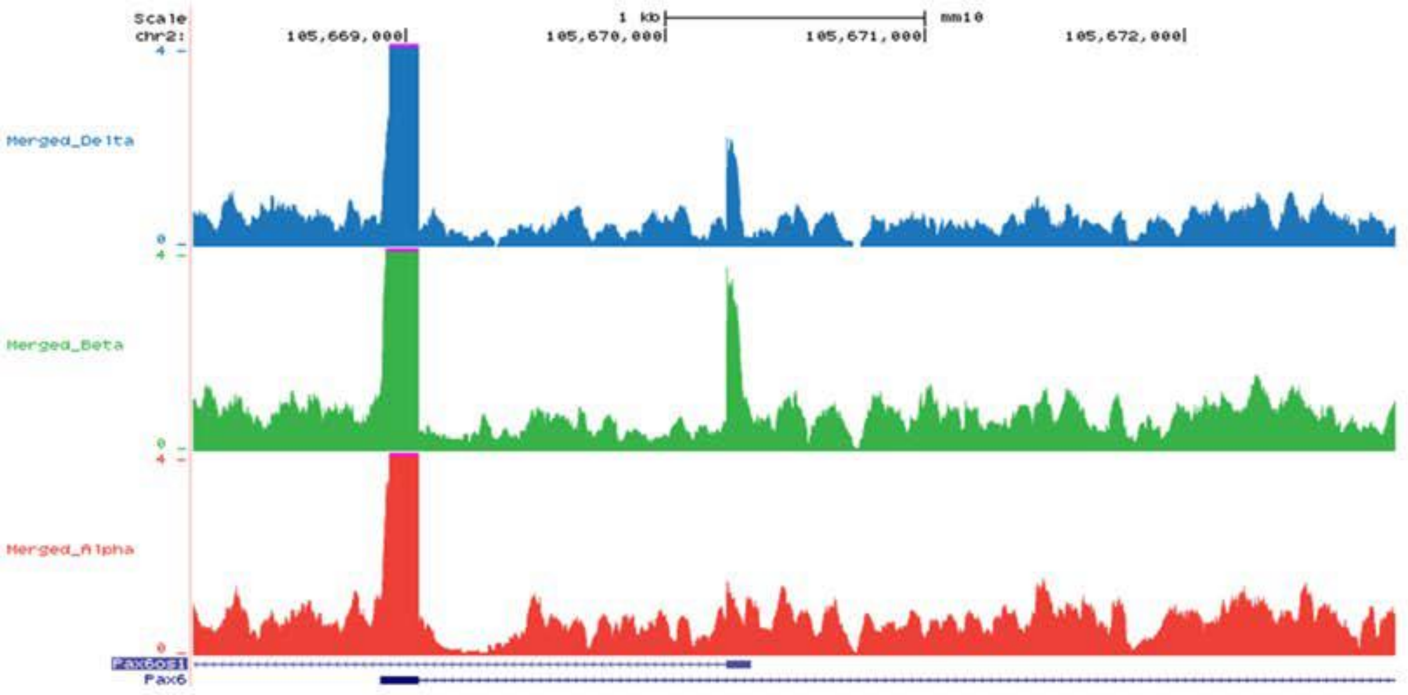


Supplemental Figure 1. Pax6os1 is mainly expressed in β-cells within pancreatic islets. Genome browser tracks showing Pax6os1 expression in δ-, β- and α- cells.

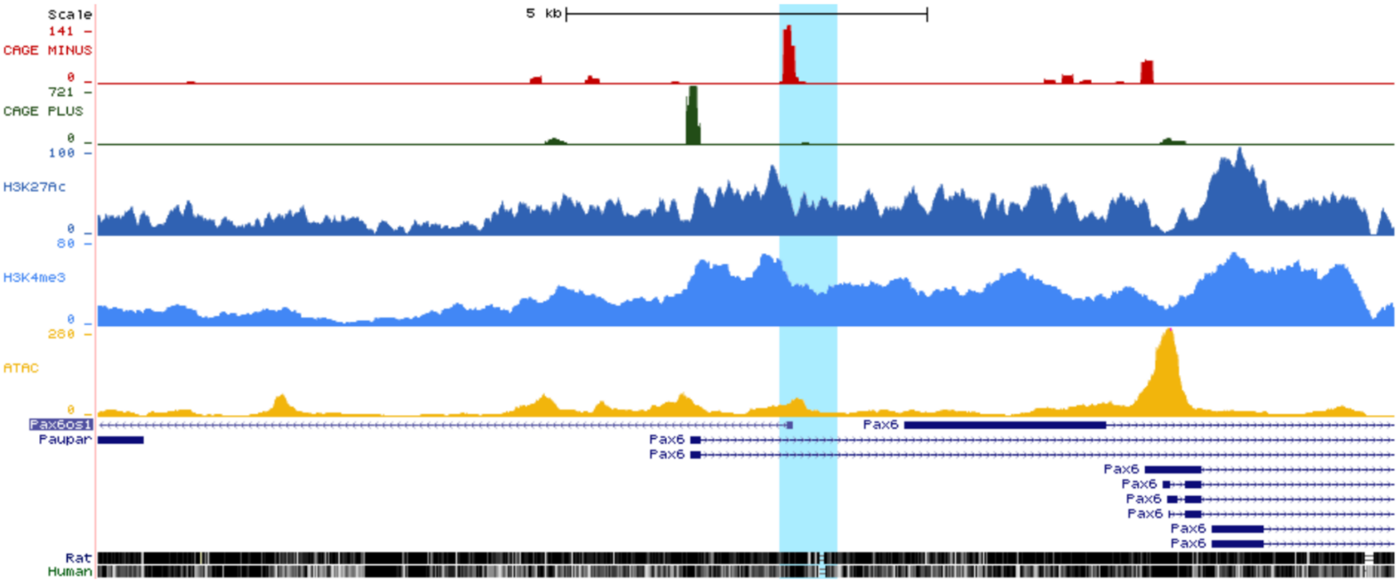


Supplemental Figure 2. Chromatin landscape of the Pax6os1 locus. Integration of ATAC-seq displaying chromatin accessibility (yellow), SLIC-CAGE showing TSSs (Minus strand, red; Plus strand, green) and H3K4me3 (Lu et al., GEO: GSE110648) and H3K27Ac (Nammo et al, E-MTAB-6719) ChIP-seq datasets (blue) in mouse islets. Annotated (GENCODE VM23) of non-coding (Pax6os1 and Paupar) and coding transcripts including spliced variants (Pax6) are shown in blue, with arrows indicating transcription directionality. The region deleted in Pax6os1 KO mice is shown in pale blue. Alignment with the human DNA track is shown at the bottom.

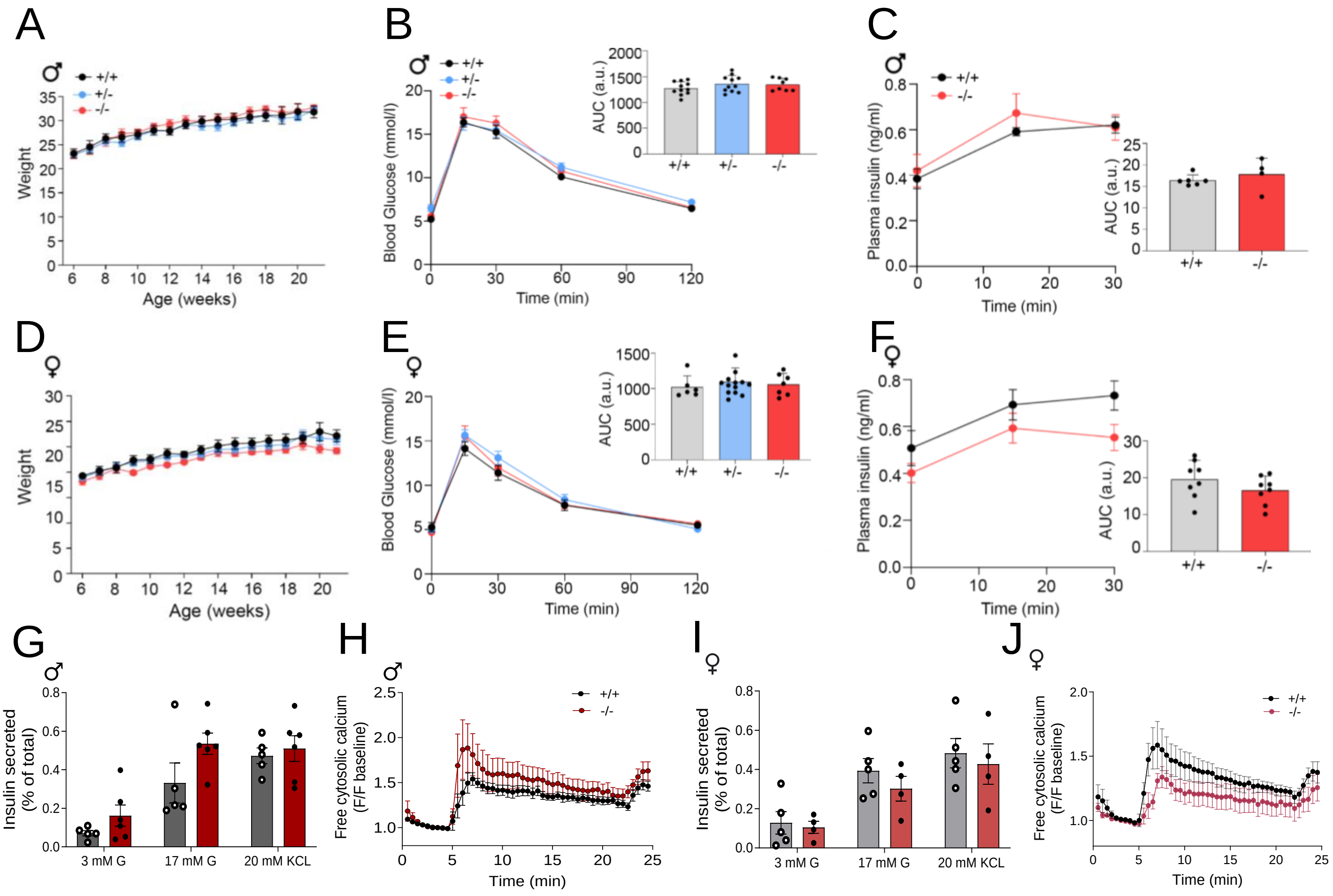


Supplemental Figure 3. Pax6os1 knockout mice display normal glucose tolerance and insulin secretion compared to WT under STD. A) Body weights (g) of wt (+/+), Pax6os1heterozygous (+/-) and Pax6os1homozygous (-/-) male mice. B) Circulating glucose levels during an intraperitoneal glucose tolerance test (IPGTT). C) Plasma insulin levels after an intraperitoneal glucose load (3g/kg). D,E,F) As in panels A, B and C in female *Pax6os1* mice. G) Insulin secreted (represented as % of the total) at different glucose concentrations and after depolarization with KCl in pancreatic islets isolated from male Pax6os1 null male mice. n= 5-6. H) Intracellular calcium in pancreatic islets isolated from male Pax6os1 null male mice. n = 3. I, J) As in panels G and H but in female mice. n= 5-4 (I); n= 3 (J). Data are represented as the mean ± SEM. *p-value < 0.05 Repeated measurements two-way ANOVA

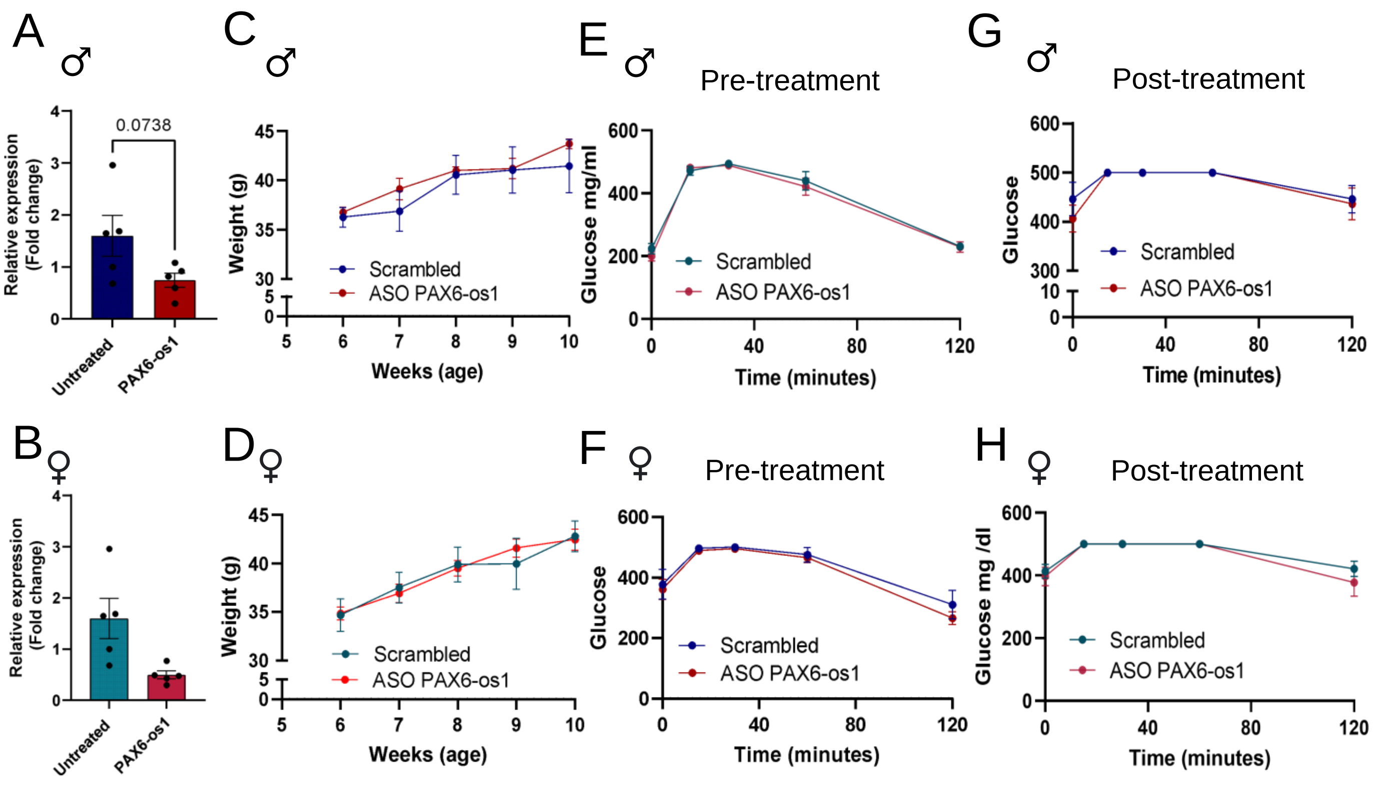


Supplemental Figure 4. Db/db mice display unaltered glucose metabolism after Pax6os1 silencing using ASO. A,B) Expression of Pax6os1 in pancreatic islets isolated from db/db mice after 4 weeks of treatment with antisense oligonucleotides targeting Pax6os1. C,D) Weights (g) od db/db male and female mice for the duration of the treatment. E,F) Glucose clearance after receiving an oral glucose load (2g/kg) in male and female mice before starting the treatment. G,H) Glucose clearance of the different experimental groups at the end of the treatment. Data are repsented as the mean ± SEM. Unpaired student t test for panels A and B and two way ANOVA repeated measurements for all the other panels, *p-value<0.05.

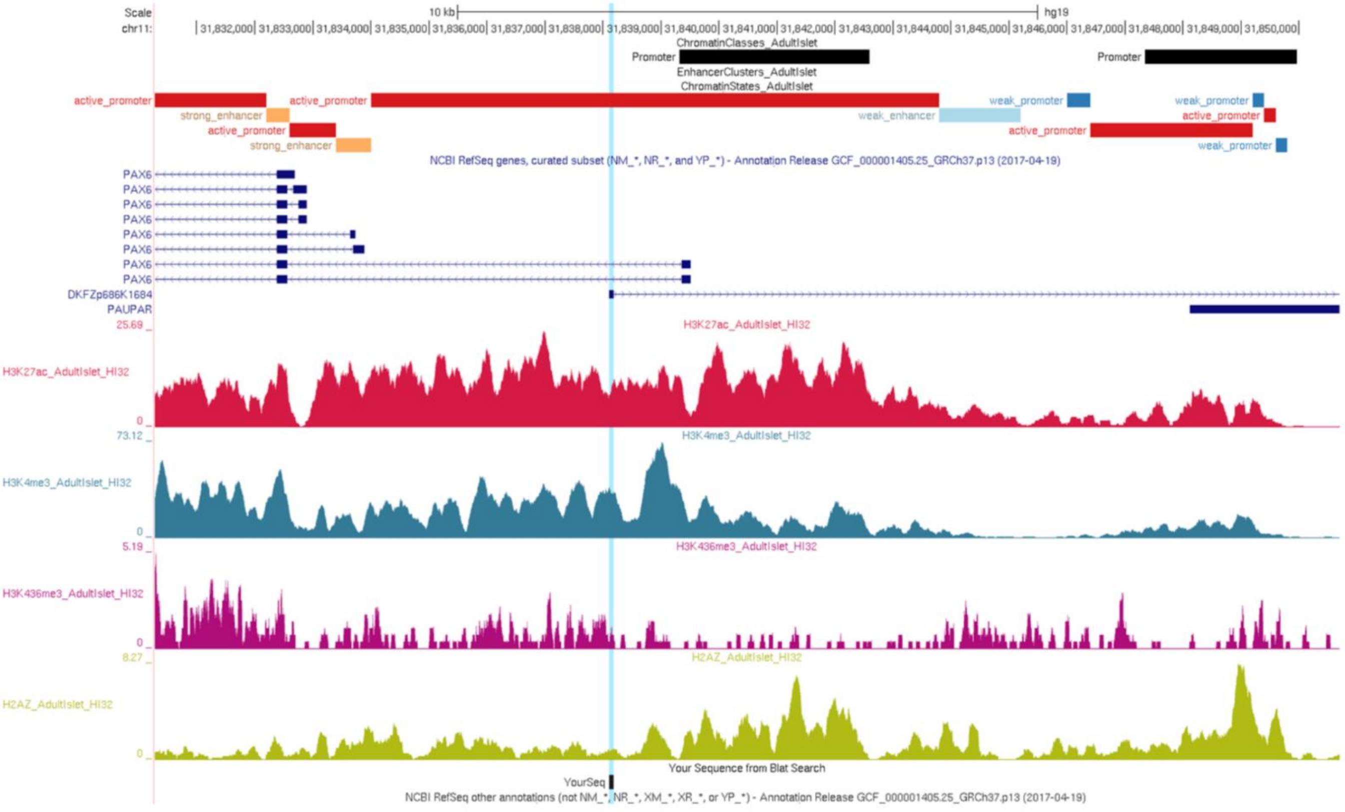


Supplemental Figure 5. Diagram showing the PAX6-AS1 deleted region in human β-cells and chromatin marks found in the PAX6-AS1 and PAX6 locus (obtained from the pancreatic islet regulome browser38. PAX6-AS1 is annotated as DKFZp686K1684
